# A common statistical misunderstanding in Psychology and Neuroscience: Do we need normally distributed independent or dependent variables for linear regression to work?

**DOI:** 10.1101/305946

**Authors:** Christian G. Habeck, Adam M. Brickman

## Abstract

Our ability to draw conclusions from experiments often relies on the application of inferential statistics, such as those included within the general-linear-modeling framework. The purpose of this commentary is to direct attention a common fallacy that can be observed in paper reviews and professional exchanges in the field of Psychology and Neuroscience pertaining to multiple regression analysis. The fallacy concerns the confusion of the requirements on the *residuals* of a linear-regression model with requirements on the *dependent or independent variables* themselves. A simple thought experiment, algebraic demonstration, and simulation study can demonstrate this misconception. It is our hope that consideration of the fallacy may help clarify distinctions between technical statistical issues and conceptual interpretation of research findings.

## Distributional demands on variables

Psychological research often utilizes inferential statistics, such as those contained in the general-linear-modeling framework. As behavioral scientists, our first exposures to inferential statistics often involve the internalization of various technical rules and potential “red flags” for interpretation of statistical results. The assumption of “normality” is often evoked as among the more important statistical assumptions. Indeed, many measurable attributes fall along a Gaussian distribution. Frequently, however, independent or dependent variables are not normally distributed. In paper reviews or scientific exchanges, we have often experienced the somewhat knee-jerk criticism that the assumptions of our inferential statistics have been violated because independent or dependent variables are not normally distributed. Apart from the exaggerated concern about the instability of ordinary-least-squares (OLS) techniques, which are ubiquitous precisely because of their very robustness against violations in their required assumptions (Pedhazur and Kerlinger 1982) and the fact that they provide unbiased, efficient and consistent estimators in most situations (Wooldridge 2002), this criticism is incorrect and confuses the requirements for proper execution of linear regression. The origin of this inappropriate criticism is not clear and we can only speculate. It might be that the tenets of normality in the distribution of variables, which are required for the correctness of *certain* parametric statistics (Micceri 1989), can all too easily be generalized to *all* statistical analyses. Textbooks of applied statistics might inadvertently contribute to this misunderstanding by not being sufficiently clear about the limited context in which normality assumptions apply. We supply a quotation of one popular text book (Altman 1991), which lays out the requirements for parametric group comparisons:

> *“[…] For independent groups, parametric methods require the observations within each group to have an approximately normal distribution, and the standard deviations in each group should be similar. If the raw data do not satisfy these conditions, a transformation may be successful. […]”*

The reader might be forgiven to take this statement as an indictment of *any* parametric technique for non-normally distributed variables, including linear regression via OLS. Formal textbook accounts of the requirements for linear regression (for instance (Cohen and Cohen 2003, Pedhazur and Kerlinger 1982,
Wooldridge 2002)) might not be as memorable as an easily generalizable caveat about normal distributions in dependent and independent variables, which is often repeated in less formal settings than peer-reviewed publications like presentations, journal reviews and internet exchanges. ^1^ Formal textbook accounts, after all, to our knowledge only list the *necessary* assumptions for linear regression to work; they usually do not list assumptions that need *not* be met.

This inappropriate insistence on normally distributed dependent or independent variables for OLS linear regression is subject of the statistical fallacy we would like to highlight. Distributional requirements, in the context of the general linear model, pertain to the distribution of residual error, *not* the distribution of independent or dependent variables. The first assumption is often referred as “constant variance” or “homoscedasticity” (Pedhazur and Kerlinger 1982). This assumption simply states that the variance of the error terms is not dependent on the value of the independent variable(s). The second assumption states that, for small samples, the errors should follow a normal distribution. If we then assume our independent variables to be known perfectly, the two aforementioned assumptions suffice to ensure that the regression-formalism produces correct results. **No distributional assumptions for dependent or independent variables themselves are required**.

We illustrate this first point with a simple thought experiment: imagine you are replicating the famous experiments by Sternberg about memory scanning (Sternberg 1966, 1969). You have a set of stimulus items and participants in the experiment are instructed to memorize them to their best ability with a subsequent test. The number of items in any of 50 trials can range from 1 to 8. We assume for simplicity that the true relationship between the number of items, *N*, and the reaction time is

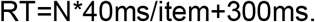

Linear regression is the obvious technique to infer both scanning speed and intercept from the observed values of *RT* and *N*. We assume the *RT* measurement is not error-free, while the number of items, *N*, is known perfectly. The true data model is thus

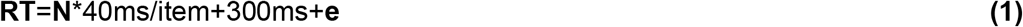

where the bolded quantities are now column vectors with 50 rows (one for each measurement), and ***e*** captures the measurement error in the dependent variable. The distribution of item numbers in ***N*** should have no influence on the scanning speed, but does it influence its empirical inference *from* the data? The common misconception assumes that non-normal distribution of the independent variable ***N*** renders the linear regression incorrect. This is wrong; instead, there is a legitimate requirement of normality on the residuals ***e***. Further, ***e*** should manifest no correlation with the independent variable ***N***.

First, we could stop and think for a moment about the influence of the distribution of item numbers on the above data model. Consider an extreme scenario without any measurement noise at all, i.e. the residuals ***e*** are vanishing. As soon as we observe the reaction time for two or more different item numbers, we can determine slope and intercept of the linear relationship exactly. We just need to solve a system of 2 equations, any further item numbers just lead to equations that are linearly dependent and thus do not add any information. This example shows that the distribution of item numbers for which the reaction time is measured is irrelevant for vanishing noise. It simply does not matter how we pick the item numbers for which we want to measure reaction times, since two values are all we need to completely determine slope and intercept. We can of course perform linear regression and obtain the correct slope and intercept estimates, but we would only need 2 observations for different item numbers. Now, we can start introduce non-vanishing noise levels and increase the amplitude of the residuals. The linear dependence of equations for different item numbers, indeed for any observation, is now absent, and this time we need as many observations as possible, but is the distribution of item numbers now important? Without the exact linear algebra, we can already venture that such a discontinuity or dependence on the level of noise would be implausible and surprising indeed: at what point would the level of noise be sufficiently small that the dependence on the distribution of item numbers disappear? Further, the very idea that probing a linear mechanism would give incorrect results if the probes were not scattered around an arbitrary mean in a Gaussian fashion strikes us as very odd indeed. It makes sense that increasing the range and number of probes would lead to more precise estimates, but why would the *correctness of the inference* be compromised?

If we now consult the linear-algebra formalism for linear regression, one can see that in the absence of a correlation between the independent variables and the residuals, the formula for the regression weight suffers no consequences. We recall that the regression weight is obtained through ordinary least-squares (Cohen and Cohen 2003). To increase mathematical convenience, we can write both item numbers and intercept term in a design matrix ***X*** as

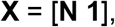

where ***N*** indicates column vector with each row denoting a different measurement, and ***1*** just assigns a value of 1 to each of the 50 rows. The data model can now be re-written as

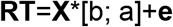

and the formula for OLS regression now proceeds according to

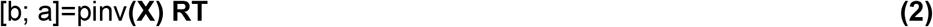

where *pinv**(X)*** indicates the Moore-Penrose pseudo-inverse (Strang 2006) of ***X***, and *b* and a, denote the slope and intercept, respectively. Simple matrix algebra shows that for the multiplication from the left with *pinv**(X)***, equation (1) reduces to equation (2) if the residuals and independent variables are uncorrelated,

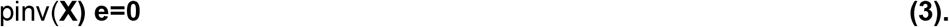

The distribution of observations in ***X*** has no effect on the linear algebra, but will the formula give the right answer for *a* and *b?* We can answer this question with a simple Monte-Carlo simulation. We use the data model given in equation (1) and sample our residuals from a normal distribution *N(0,(10ms)^2^)*. For the item numbers, we sample from a skewed distribution for which the probability of an item number goes down linearly with the item number itself. We perform linear regression and can see how close the inferred answers for *a* and *b* come to the true answers of 300ms and 40ms /item, respectively. Does it matter that the distribution of values in *N* departs from normality? Next we perform the same simulations, but this time we sample from a distribution of residuals that violates the assumption of normality, as it is both biased and non-Gaussian. The results can be seen in Figure 1 below.

**Figure 1:**
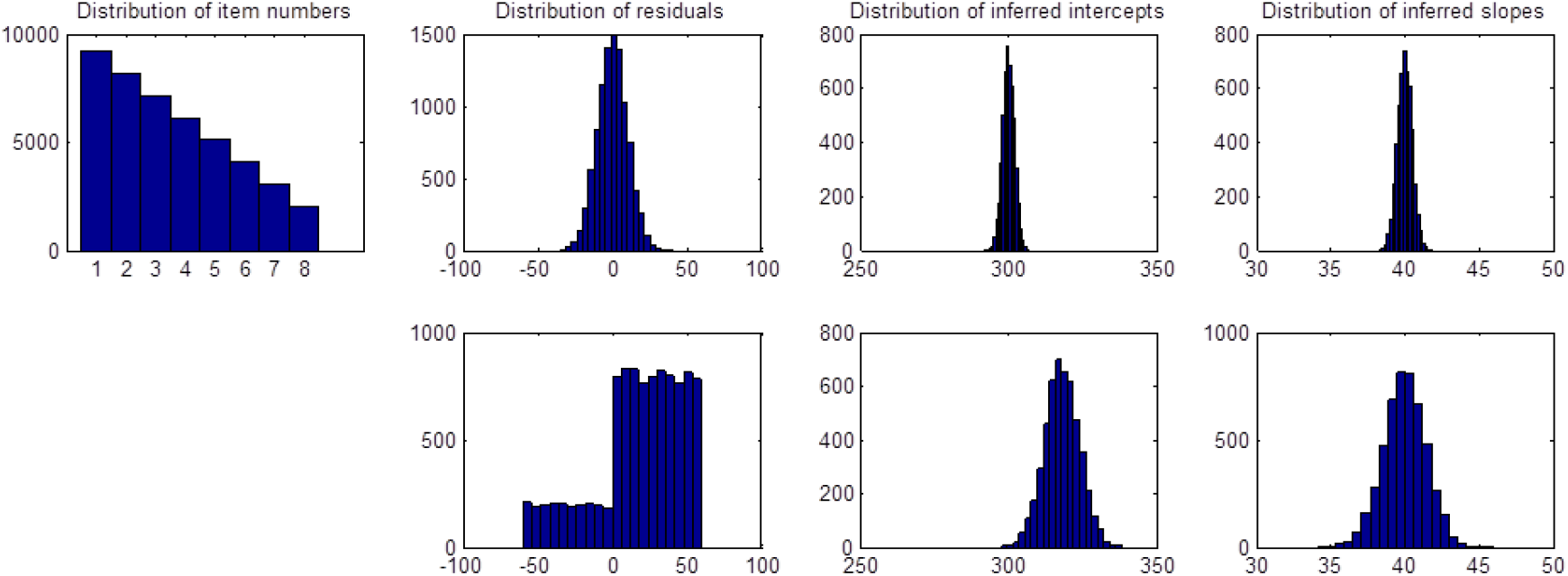
Impact of non-normal independent variables and residuals on the correctness of linear regression results. Two Monte-Carlo simulations with 10,000 iterations for 50 observations were conducted. The left most upper panel shows the distribution of item numbers from which the independent variables were sampled for both simulations. The remaining three panels in both top and bottom row show the distribution of residuals used in the data model, and the empirical distribution of inferred intercepts and slopes from conducting the regressions. The top row features the case of Normally distributed residuals. The bottom row, on the other hand, shows the case of ill-behaved residuals. The inferred intercepts are biased and more variable than in well-behaved case; the slopes are unbiased but more variable. In both cases, however, parametric variability estimates are in line with empirical reality, i.e. the means of the parametric error estimates are equal to the empirical standard deviations of observed slopes and intercepts.

The most left upper panel shows the clearly non-normal distribution of item numbers from which the independent variables were sampled for both simulations. The remaining three panels in both top and bottom row show the distribution of residuals used in the data model, and the empirical distribution of inferred intercepts and slopes from conducting the regressions. The top row features the case of normally distributed residuals. The bottom row, on the other hand, shows the case of ill-behaved residuals. The inferred intercepts are biased and more variable than in well behaved case; the slopes are unbiased, but also more variable. In both cases, however, parametric variability estimates are in line with empirical reality, i.e. the means of the parametric error estimates are equal to the empirical standard deviations of observed slopes and intercepts (see caption of Figure 1). Thus, for the non-normally distributed residuals, parametric and empirical estimates are equal – the obtained intercept value would just be shifted from its true underlying value for both techniques. We will outline some simple diagnostic measures that would flag this case clearly.

These results show that, in line with our simulation results and the general requirements for OLS linear regression, **regardless of the distribution of independent variables, the parametric formalism for linear regression works well if the residuals are well behaved**. The formalism does not work well when the distribution of the residuals violates the requirements of normality. This latter case is rare. From our anecdotal experience the formalism is quite robust, even when the residuals are not Gaussian. Theoretically, we state the Gauss-Markov Theorem (Pedhazur and Kerlinger 1982, Wooldridge 2002): when errors are unbiased and uncorrelated with independent variables and the size of the error is independent of the magnitude of any independent variable, the least-squares formalism always leads to the best linear unbiased estimate (BLUE), i. e. the estimates coming out of the regression are always unbiased and always have the minimal possible variance, regardless of the distribution of errors.

We showed that the distribution of the independent variables is irrelevant if the assumptions of homoscedasticity and normality are fulfilled for the residuals. But what about the distribution of the *dependent* variable? Anecdotally, the distribution of the dependent variable is often the bigger concern and one can occasionally hear the rule of thumb “Normality is not so important for independent variables, but very important for dependent variables”. We can easily demonstrate that this declaration is also untrue, consistent with the requirements laid out in formal textbook accounts. Non-normally distributed dependent variables are hard to contrive for situations where there is a real effect in the data and the residuals are normally distributed. In our previous examples, regardless of the distribution of the independent variables, the dependent variables often end up looking normally distributed. This is not surprising, since for normally distributed residuals ***e*** that are not correlated with the independent variable, the mean and variance for the Sternberg model is derived by the standard formula for error propagation as

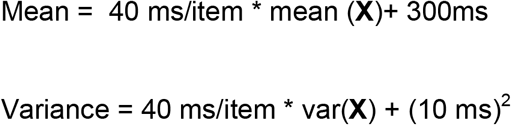

The larger the noise levels in the residuals, the more the shape of the distribution for the dependent variable looks like a Gaussian, as shown in Figure 2 below.

**Figure 2:**
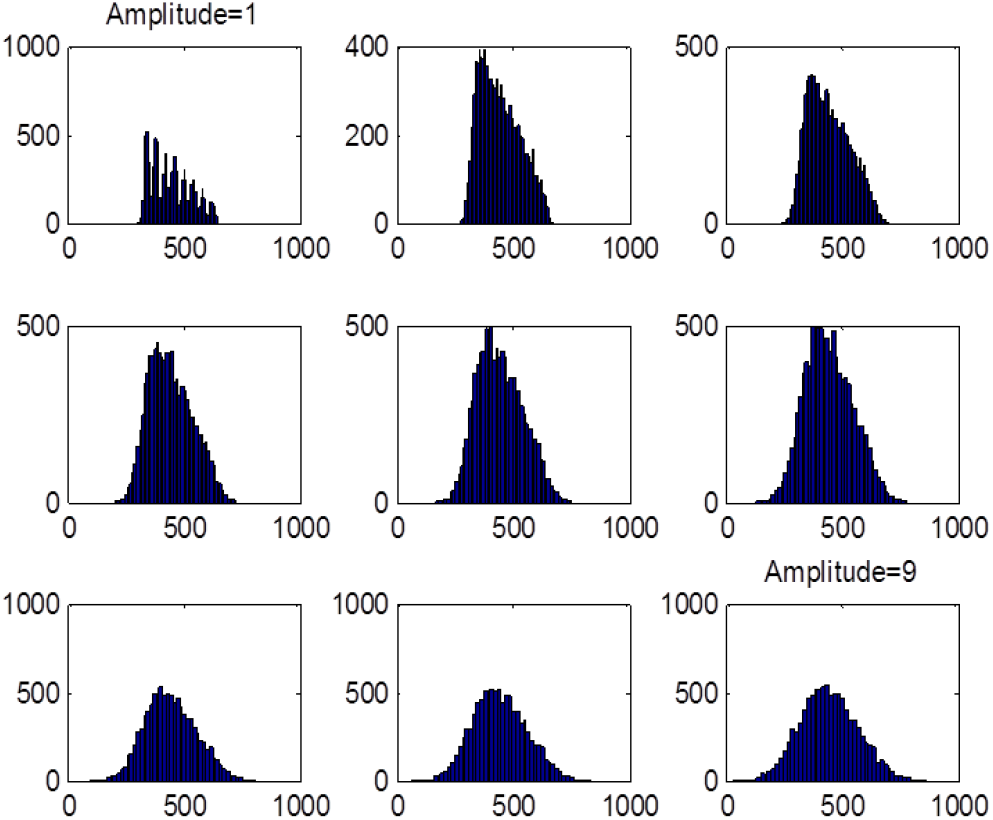
Influence of noise level on the shape of the distribution of the dependent variable. 10,000 values were generated according for the RT-Sternberg model, using the distributions of independent variables and residuals according to the top row of Figure 1. The residuals’ amplitudes were scaled up with factor that ranged from 1 to 9, going from left to right, and top to bottom in the matrix of scatter plots. With increasing noise levels, the distribution of the dependent variable becomes Gaussian.

For a real effect in the data, dependent variables might often follow a Gaussian distribution, but this does not in any way imply that Gaussianity of dependent variables is *necessary* for linear regression to work. We can show this for null data where independent variables and residuals conform to the distributions shown in the top row of Figure 1, but the dependent variable is randomly sampled from the uniform distribution U(0,1). We can compare parametric p-levels and p-levels obtained from a permutation test with 10,000 iterations for the inferred slope parameter. If non-Gaussian dependent variables were to cause inflation of p-levels, the parametric p-levels should be consistently lower than the empirical p-levels obtained with the permutation test. Figure 3 shows the comparison of both types of p-levels for 500 null scenarios.

**Figure 3:**
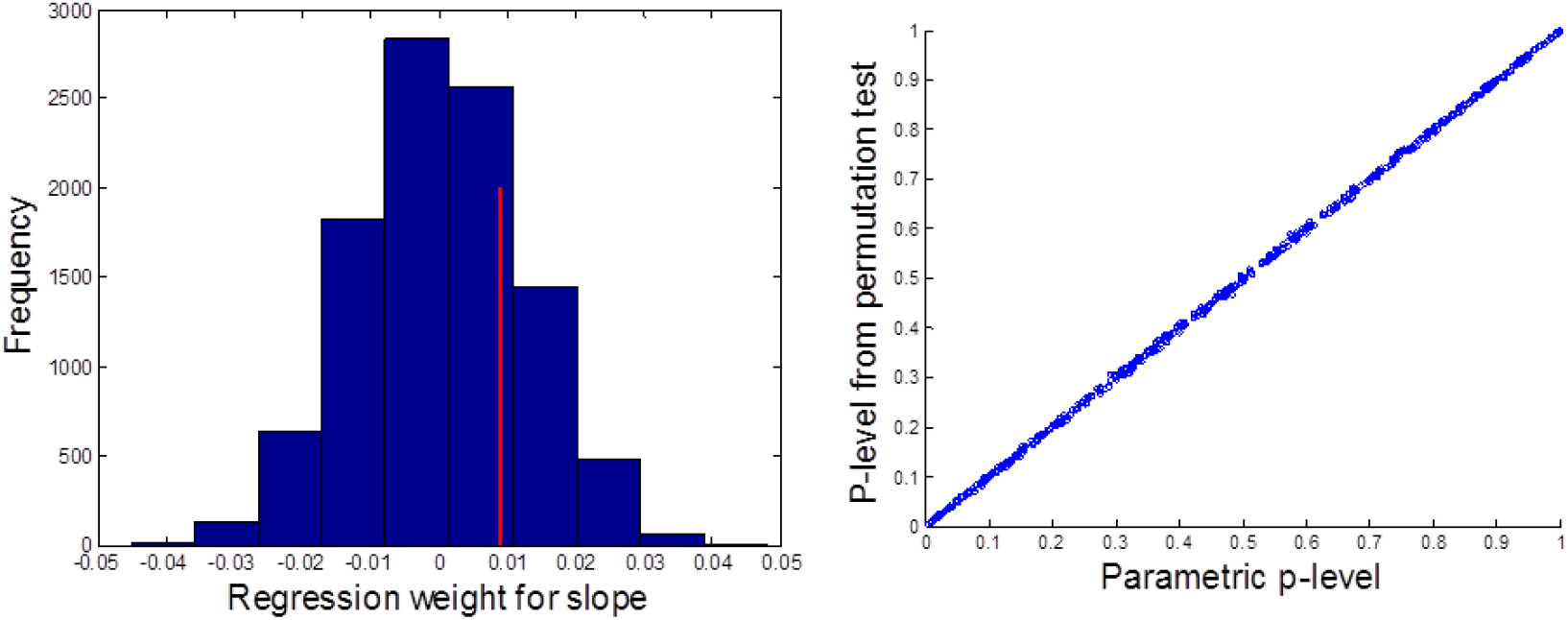
Investigation of possible inflation of false positives for non-normal dependent variables under null effects. Each time independent variables and residuals were sampled from the skewed and Normal distributions, respectively, used in the top row of Figure 1, while the dependent variable was randomly sampled from the uniform distribution U(0,1). Parametric p-levels were compared to p-levels generated from permutation tests with 10,000 iterations for 500 null scenarios. One example is shown in the left panel: the null distribution of inferred slope values is shown with the point estimate marked with the red vertical line. The two-tailed p-level obtained from the permutation test was 0.4834 while the parametrically estimated value was 0.4813. The right panel shows a scatter plot of parametric p-levels against p-levels obtained from permutation tests for all 500 simulations: one can appreciate that there is no discrepancy, i.e. the results fall on the unity line (rather than sloping upwards above the unity line), and non-normal dependent variables do not cause false positives when treated with standard regression techniques.

Figure 3 shows the results of a permutation test for a randomly picked example and a scatter plot of both types of p-levels for all 500 examples. The figure shows that the parametric-regression formalism works just as well for non-Gaussian dependent variables and does not lead to an under-estimation of false positives.

For most experimental situations, worries about distributions, even in the residuals, are thus unnecessary.More important are “specification errors” (Pedhazur and Kerlinger 1982), which lead to collinearities between residual errors and independent variables; such collinearities violate the fundamental assumption of the Gauss-Markov theorem, invalidating both BLUE property and parametric estimates, and possibly estimated type-I-error probabilities.

What should one do in case of suspicion that the residuals are not well behaved, leaving to incorrect p-values and possibly biased regression weights? We can give some practical recommendations for easy checks that validity requirements are met and techniques that do not rely on formal assumptions. There are no absolute thresholds or cutoffs, but researchers could check the model residuals in the following ways (Pedhazur and Kerlinger 1982, Cohen and Cohen 2003). We are slightly conflicted whether to issue such a recipe for checking at all since it might cause undue worry. In our anecdotal experience in 99% of the cases one does not have to be concerned about residuals at all and can safely forget about them:

1. Plot the residuals as a histogram, similarly to Figure 1 and visually inspect the distribution for an obvious bias, skewness and kurtosis.
2. Check the orthogonality of residuals to all independent variables, i.e. compute the correlation of each predictor with the residuals and determine whether a significant correlation is present.
3. Plot the residuals against the independent variables – is any kind of structure or relationship visible, even though nominal correlation values are low?

In addition to these diagnostic measures, researchers can always perform a bootstrap-resampling procedure (Efron 1982, Efron and Tibshirani 1993) and resample from the dependent and independent variables with replacement at least several thousand times, each time keeping the assignment of dependent and independent variables intact and performing the linear regression in Equation (2). This approach produces an empirical distribution for the obtained intercept and slope, and obviates any reliance on parametric formulae or particular assumptions of independence between residuals and independent variables.

We return to the main goal of clarification in this technical note and state our summary conclusion again: **if the residuals in a linear regression are well behaved, the distribution of dependent or independent variables is irrelevant**.

Transforming either variable, for instance using logarithms or other transcendental functions, makes no difference to the correctness of the statistical inference. Does this mean one should never transform variables? – No, since there might be good reasons for variable transformations due to (1) interpretability of the results, or (2) non-statistical prior information about the correct model. For (1), a prime example is the choice of logistic, over linear, regression. Predicting logit-functions as outcomes leads to a bounded sigmoidal relationship between an independent variable and a class probability, which makes more intuitive sense than an unbounded linear relationship between the independent variable and a, somewhat arbitrary, class label. For (2), we can refer to our earlier example of the Sternberg memory-scanning experiment. On statistical grounds, we would be entirely justified to transform the dependent variable (= reaction time) by taking the natural logarithm with no consequence for the correctness of the statistical inference; however, this is clearly not the right thing to do since we would be modeling an incorrect *exponential*, rather than *linear*, relationship between the number of memory items and reaction time. Our practical advice for researchers is that for linear regression, **transforming variables might be appropriate on account of interpretability or prior model-constraints, but not on account of the distribution of independent or dependent variables. For linear regression, such considerations of distributional “hygiene” should <u>not</u> play any role in motivating variable transformations**.

## Conclusion

We think the fallacy we pointed out in this commentary is remedied easily since formal textbook accounts are already completely clear about the requirements of linear regression. Given that the fallacy is due to a simple confusion of residuals with the independent or dependent variables, it is imperative that researchers try to eliminate it from professional exchanges. In our anecdotal experience the fallacy can unnecessarily take up considerable effort for journal editors, reviewers and authors – effort that would be better spent on more pressing issues of science and experimental design.

## Acknowledgments

We thank Yian Gu, Yunglin Gazes, and Tamer Abdelgawad for a careful reading of the manuscript with many helpful suggestions.

## Footnotes

1 Examples for consulting exchanges about the question of the distribution of variables can be found in on the web sites below: http://www.researchgate.net/post/Do_we_need_normal_distribution_of_dependent_variable_when_working_with_ordinary_least_squares_or_other_linear_regression_method http://www.angelfire.com/wv/bwhomedir/notes/normality_and_regression.txt

